# Single-cell transcriptomics reveals expansion of cytotoxic CD4 T-cells in supercentenarians

**DOI:** 10.1101/643528

**Authors:** Kosuke Hashimoto, Tsukasa Kouno, Tomokatsu Ikawa, Norihito Hayatsu, Yurina Miyajima, Haruka Yabukami, Tommy Terooatea, Takashi Sasaki, Takahiro Suzuki, Matthew Valentine, Giovanni Pascarella, Yasushi Okazaki, Harukazu Suzuki, Jay W. Shin, Aki Minoda, Ichiro Taniuchi, Hideyuki Okano, Yasumichi Arai, Nobuyoshi Hirose, Piero Carninci

## Abstract

Supercentenarians, people who have reached 110 years of age, are a great model of healthy aging. Their characteristics of delayed onset of age-related diseases and compression of morbidity imply that their immune system remains functional. Here we performed single-cell transcriptome analysis of 61,202 peripheral blood mononuclear cells (PBMCs), derived from seven supercentenarians and five younger controls. We identified a marked increase of cytotoxic CD4 T-cells (CD4 CTLs) coupled with a substantial reduction of B-cells as a novel signature of supercentenarians. Furthermore, single-cell T-cell receptor sequencing of two supercentenarians revealed that CD4 CTLs had accumulated through massive clonal expansion, with the most frequent clonotypes accounting for 15% to 35% of the entire CD4 T-cell population. The CD4 CTLs exhibited substantial heterogeneity in their degree of cytotoxicity as well as a nearly identical transcriptome to that of CD8 CTLs. This indicates that CD4 CTLs utilize the transcriptional program of the CD8 lineage while retaining CD4 expression. Our study reveals that supercentenarians have unique characteristics in their circulating lymphocytes, which may represent an essential adaptation to achieve exceptional longevity by sustaining immune responses to infections and diseases.

**Significance:** Exceptionally long-lived people such as supercentenarians tend to spend their entire lives in good health, implying that their immune system remains active to protect against infections and tumors. However, their immunological condition has been largely unexplored. We profiled thousands of circulating immune cells from supercentenarians at single-cell resolution, and identified a large number of CD4 T-cells that have cytotoxic features. This characteristic is very unique to supercentenarians, because generally CD4 T-cells have helper, but not cytotoxic, functions under physiological conditions. We further profiled their T-cell receptors, and revealed that the cytotoxic CD4 T-cells were accumulated through clonal expansion. The conversion of helper CD4 T-cells to a cytotoxic variety might be an adaptation to the late stage of aging.

---

Supercentenarians are rare individuals who reach 110 years of age. They are endowed with high resistance to lethal diseases such as cancer, stroke, and cardiovascular disease (1–4). Demographers in Canada estimated that the chance of living more than 110 years is as low as 1 in 100,000 (http://www.forum.umontreal.ca/forum_express/pages_a/demo.htm). According to the population census covering the whole territory of Japan in 2015 (http://www.stat.go.jp/english/data/kokusei/2015/pdf/outline.pdf), the number of centenarians was 61,763, of which only 146 were supercentenarians. A distinctive feature of supercentenarians is a long healthy life-span, maintaining relatively high cognitive function and physical independence even after 100 years of age (5, 6). In other words, many supercentenarians can spend almost their entire lives in good health due to the delayed onset of age-related diseases and compression of morbidity (7). Therefore, supercentenarians can be considered a good model of successful aging, and understanding their attributes would be beneficial for super-aging societies. Many functions of the immune system show a progressive decline with age, a phenomenon known as immunosenescence, leading to a higher risk of infection, cancer, and autoimmune diseases (8, 9). A low level of inflammation is the best predictor of successful aging at extreme old age, indicating the importance of maintaining the immune system (10). Age-related alterations are apparent in two primary lymphoid organs, thymus and bone marrow, which are responsible for the development of mature lymphocytes (11). In particular, elderly hematopoietic stem cells in bone marrow exhibit a myeloid-biased differentiation potential (12, 13), which causes changes in the cell population of peripheral blood.

Numerous studies have examined age-related alterations in whole blood and peripheral blood mononuclear cells (PBMCs), derived from healthy donors in a wide range of age groups. Fluorescence activated cell sorting (FACS) and transcriptome sequencing technologies, which are extensively used to profile circulating immune cells, have revealed that the population makeup and expression levels of peripheral lymphocytes change dynamically with age. For example, the absolute number and percentage of peripheral blood CD19 B-cells decrease with age (14–16). Naïve T-cell numbers tend to decrease according to age, whereas antigen-experienced memory T-cell numbers increase with concomitant loss of co-stimulation factors CD27 and CD28 (17). This tendency is more pronounced for CD8 T-cells in cytomegalovirus seropositive donors (18). In parallel, transcriptome studies have reported a large number of age-associated genes in bulk peripheral blood that can be used to predict ‘transcriptomic age’ (19). However, most of the studies targeted donors from young to 100 years old, and the circulating immune cells in supercentenarians remain largely unexplored.

Single-cell transcriptomic methods have rapidly evolved in recent years. The accuracy of quantifying gene expression and the number of cells captured per experiment have been dramatically improved (20, 21). These methods have been applied to various subjects such as finding signatures of aging in the human pancreas (22), observing infiltrating T-cells in tumors (23, 24), and characterizing diversity of cell types during brain development (25). Here we profiled circulating immune cells in supercentenarians at single-cell resolution and identified unique signatures in supercentenarians that could characterize healthy aging.

## Results

### Single-cell transcriptome profiling of PBMCs

We profiled fresh PBMCs derived from seven supercentenarians (SC1–SC7) and five controls (CT1–CT5, aged in their 50s to 80s) by using droplet-based single-cell RNA sequencing technology (10X Genomics) (26) (Figs. 1a and S1a). The total number of recovered cells was 61,202 comprising 41,208 cells for supercentenarians (mean: 5,887 cells) and 19,994 cells for controls (mean: 3,999 cells), which is in the normal range of median gene and UMI counts per cell reported in the 10XQC database (http://10xqc.com/index.html) (Figs. 1b and S1b). Based on their expression profiles, we visualized the cells in two-dimensional space using t-SNE (t-distributed stochastic neighbor embedding), a method for non-linear dimensionality reduction. Using a k-means clustering algorithm, we found ten distinct clusters representing different cell types (Figs. 1c and S1c). We identified the major cell types comprising PBMCs, including: T cells (TC1 and TC2 clusters) characterized by *CD3* and T-cell receptor (*TRAC*) expression, B cells (BC cluster) characterized by *MS4A1* (*CD20*) and *CD19* expression, natural killer cells (NK cluster) characterized by *KLRF1* expression, two subsets of monocytes (M14 and M16 clusters) characterized *CD14* and *FCGR3A* (*CD16*) expression, respectively, and erythrocytes (EC cluster) characterized by *HBA1* (hemoglobin alpha locus 1) expression (Figs. 1d and S1d). We also found three small clusters, annotated as MKI67+ proliferating cells (MKI), dendritic cells (DC), and megakaryocytes (MGK), based on the expression of established marker genes (Fig. S1e). Each of the clusters consisted of cells from more than eleven different donors, and there was no obvious batch effect leading to library-specific clusters (Fig. S1c).

**Figure 1.**
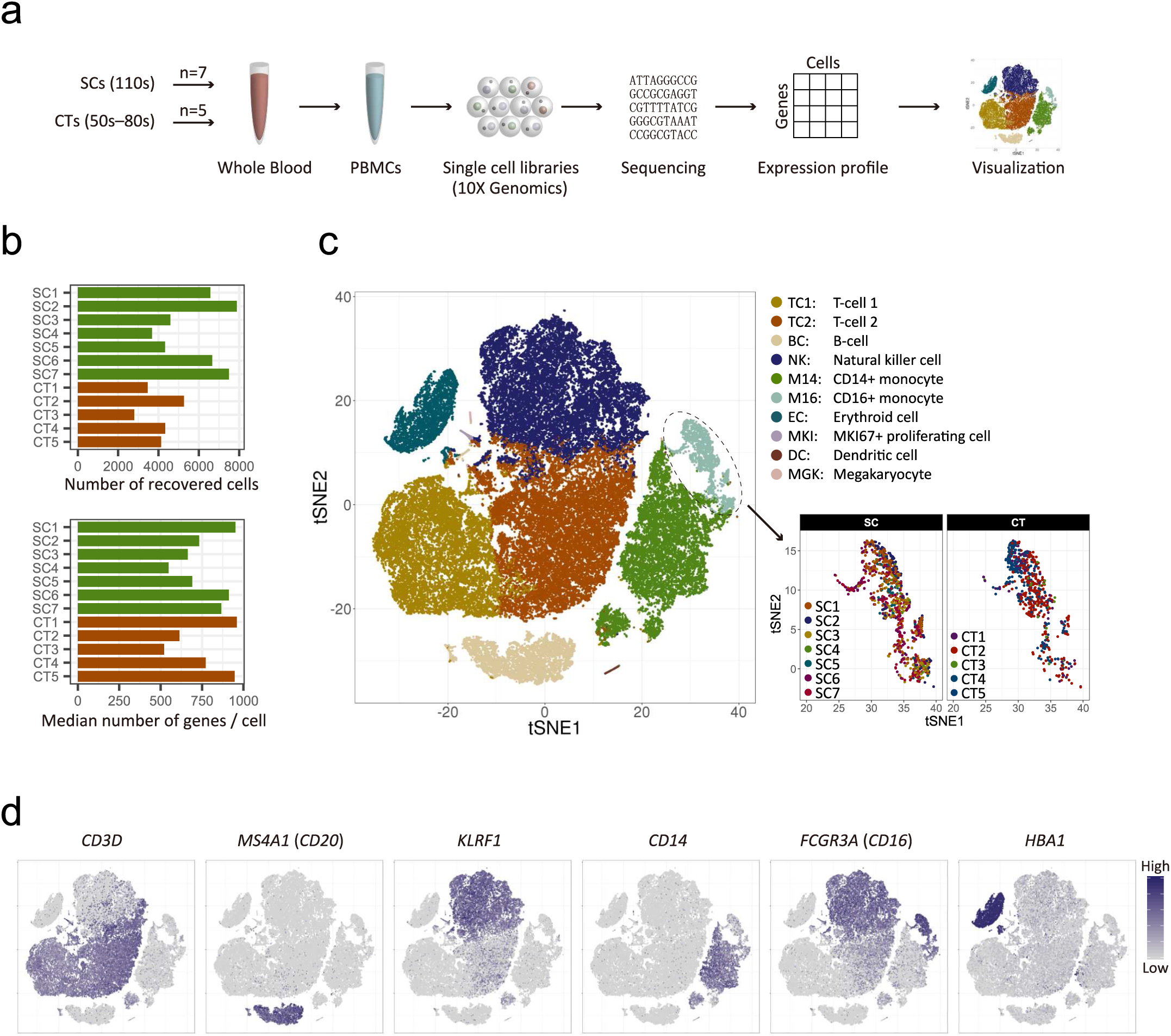
Single-cell transcriptome profiling of PBMCs of supercentenarians and controls. **a.** Schematic representation of single-cell transcriptome experiments, from blood sample collection to visualization. **b.** The number of recovered cells that passed quality control, and the median number of genes per cell for each of the donors (seven supercentenarians, SC1−SC7; and five controls, CT1–CT5). **c.** Two-dimensional tSNE visualization of 61,202 PBMCs. Different colors in the main map represent ten clusters (cell types) defined by the k-means clustering algorithm. Different colors in the enlarged view represent the 12 donors, separated into supercentenarians (left) and controls (right). **d.** Expression of marker genes for six major cell types; cell positions are from the tSNE plot in **c**.

### Significant reduction of B cells

In previous FACS analyses using cell-surface markers, various age-associated population changes were observed in human PBMCs, such as B-cell reduction (15) and loss of naïve CD8 T-cells (18). To understand whether supercentenarians follow the common population changes, we compared the percentages of the immune cells in PBMCs between the supercentenarians and controls. Among the identified cell types in our single-cell transcriptome analysis, B cell numbers were significantly decreased in the supercentenarians compared with the controls (*P* = 0.0025, Wilcoxon rank sum test) (Fig. 2a). The median percentage of B cells in the seven supercentenarians (2%) was far below that in the controls (11%) and the reference values reported in a previous cohort study (27); in contrast, the populations of the other cell types were relatively stable and did not significantly change compared with the controls (Figs. 2a and S2a). The reduction of B cells was validated by FACS analysis of four supercentenarians (SC1–SC4) and three controls (CT1–CT3), which showed low levels of CD3− and CD19+ B-cell populations in supercentenarians (Figs. 2b and S2b). We also confirmed that the percentages of major cell types (B cells, T cells, natural killer cells, and CD14+ monocytes) in PBMCs were consistent with those measured by FACS using canonical markers (Figs. 2c and S2b). We further clustered the B cells into three distinct subtypes (BC1, BC2, and BC3) by using k-means clustering (Figs. 2d and S2c). BC1 corresponds to naïve B-cells due to the presence of *IGHD*, an immunoglobulin isotype expressed before class switching, and absence of the activation marker *CD27*. BC2 corresponds to quiescent memory B-cells, characterized by expression of *CD27*, *IGHG1*, and *IGHA1* (Figs. 2e and S2d). BC3, which accounts for a small fraction, albeit one with contributions from all donors, shows distinct features of plasma cells such as high levels of immunoglobulins (*IGHA* and *IGHG*), expression of *CD38*, and loss of *MS4A1* (*CD20*) (Figs. 2e and S2e). Among these three B-cell subtypes in PBMCs, the percentage of naïve B-cells was significantly lower in supercentenarians compared with the controls (*P* = 0.005, Wilcoxon rank sum test) and the percentage of memory B-cells also tended to be lower in supercentenarians but the difference was not significant (*P* = 0.073) (Fig. 2f). Therefore, the decrease in naïve B-cells was the major source of the B-cell reduction in supercentenarians.

**Figure 2.**
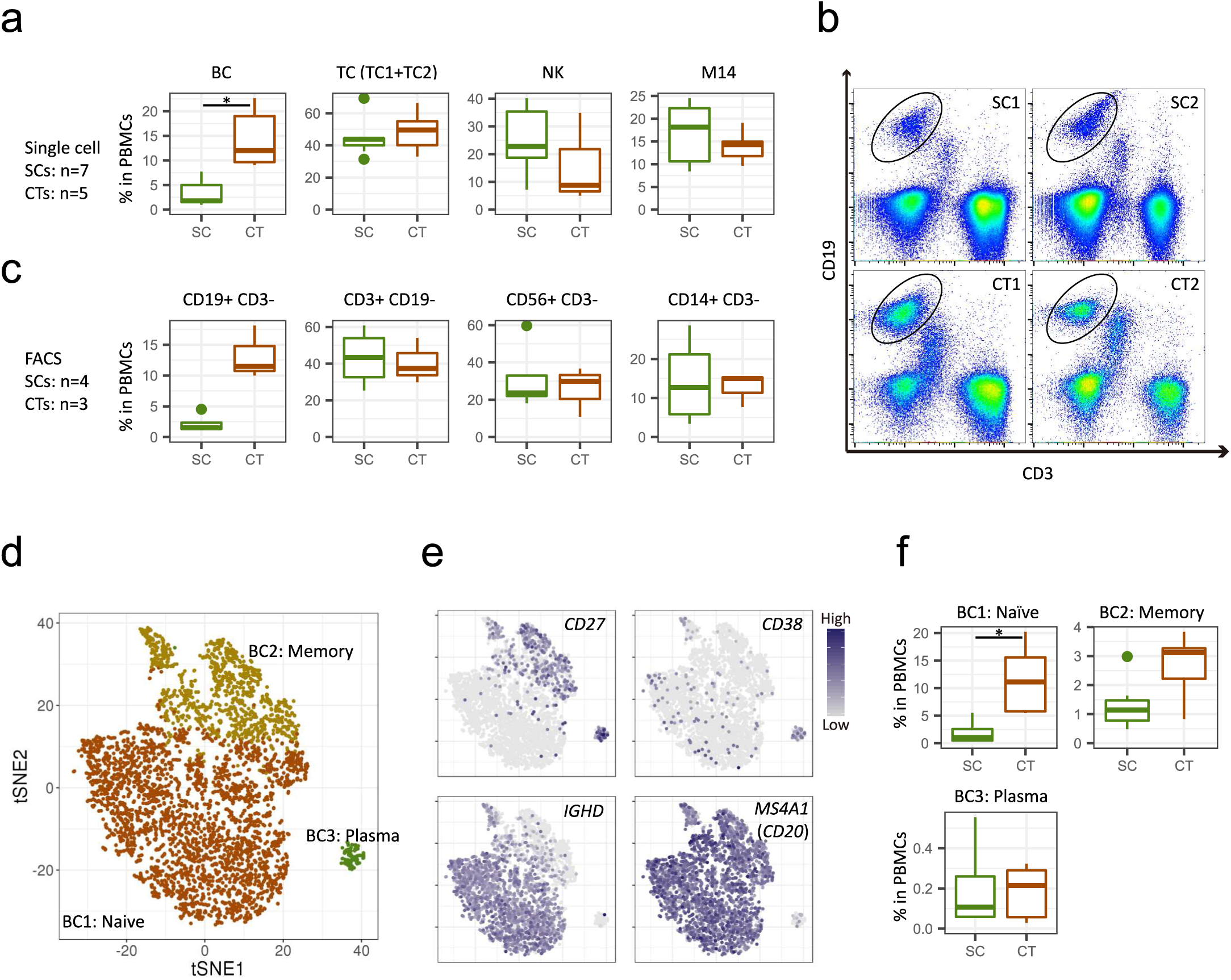
Significant reduction of B cells in supercentenarians. **a.** Boxplots of the percentage of each cell type (defined by single-cell RNA-Seq) in PBMCs of seven supercentenarians (SC1– SC7) and five controls (CT1–CT5)—the boxes extend from the 25th to 75th percentile and encompass the median (horizontal line). BC, B cell; TC, T cell; NK, natural killer cell; M14, CD14+ monocyte. *, *P* < 0.05 (Wilcoxon rank sum test); no asterisk means not significant. **b.** Representative FACS plots showing CD19+ B-cells; the plots for other donors are shown in Fig. S2b. **c.** Boxplots of the percentage of each cell type (defined by FACS) in PBMCs of four supercentenarians SC1–SC4 and three controls CT1–CT3. **d.** Two-dimensional tSNE visualization of B cells from all 12 donors. Different colors represent the three clusters defined by the k-means clustering algorithm. **e.** Expression of the indicated markers for B-cell subtypes; cell positions are from the tSNE plot in **d**. **f.** Boxplots of the percentage of each B-cell subtype (defined by k-means clustering) in PBMCs of seven supercentenarians (SC1–SC7) and five controls (CT1–CT5). *, *P* < 0.05 (Wilcoxon rank sum test); no asterisk means not significant.

### Expansion of cytotoxic T-cells in supercentenarians

In contrast to the profound reduction of B cells, the T-cell fraction remained stable at around 40% of PBMCs according to both the transcriptome data (TC in Fig. 2a) and the FACS analysis (CD3+CD19− in Fig. 2c). However, two T-cell clusters, TC1 and TC2, were imbalanced between supercentenarians and controls: TC1 was significantly diminished (*P* = 0.0025, Wilcoxon rank sum test), whereas TC2 was significantly expanded (*P* = 0.0025) in supercentenarians (Fig. 3a). To better understand this T-cell specific population shift, we extracted all the cells from TC1 and TC2 for further analysis using the Seurat R package (version 2.3.0) (28). A clustering algorithm based on shared nearest neighbor modularity optimization implemented in Seurat produced two major clusters: Seurat_TC1 and Seurat_TC2, corresponding to the original TC1 and TC2 clusters (Figs. 3b and S3a). We then compared these two clusters and identified 332 differentially expressed genes, of which the most significant gene distinctively expressed in Seurat_TC2 was *NKG7*, a component of granules in cytotoxic lymphocytes. In addition, the top 20 most significant genes included multiple genes encoding cytotoxic effector molecules responsible for the perforin/granzyme apoptosis pathway, such as *GZMH*, *GZMB*, *GZMA*, and *PRF1* (Fig. 3c and S3b). In contrast, Seurat_TC1 was characterized by expression of *CCR7* and *SELL* (encoding CD62L), which are required for lymph node migration (Fig. S3c). These genes are normally expressed in naïve and central memory T-cells, but not in cytotoxic effector memory T-cells (29), indicating that the primary factor separating the two clusters is cytotoxicity. Perforin/granzyme+ cells were predominantly found in the supercentenarians (Fig. 3d), whereas CCR7+ non-cytotoxic cells were more abundant in the controls (Fig. S3d). We then examined how many of the four cytotoxic genes (*GZMH*, *GZMB*, *GZMA*, and *PRF1*) showed detectable expression in each single cell. As expected, for both the supercentenarians and controls, the vast majority of cells in the non-cytotoxic cluster (Seurat_TC1) expressed either zero or one cytotoxic gene(s) (Fig. 3e left). In the cytotoxic cluster (Seurat_TC2), cells that expressed all four genes were abundant in supercentenarians but rare in controls, indicating that the level of cytotoxicity per cell might be higher in supercentenarians (Fig. 3e right). Cytotoxic T-cells were expanded in supercentenarians, reaching 90% of T cells in some individuals (Fig. 3f). This was in sharp contrast to controls where cytotoxic T-cells made up approximately 10% to 20% of the total T-cell population.

**Figure 3.**
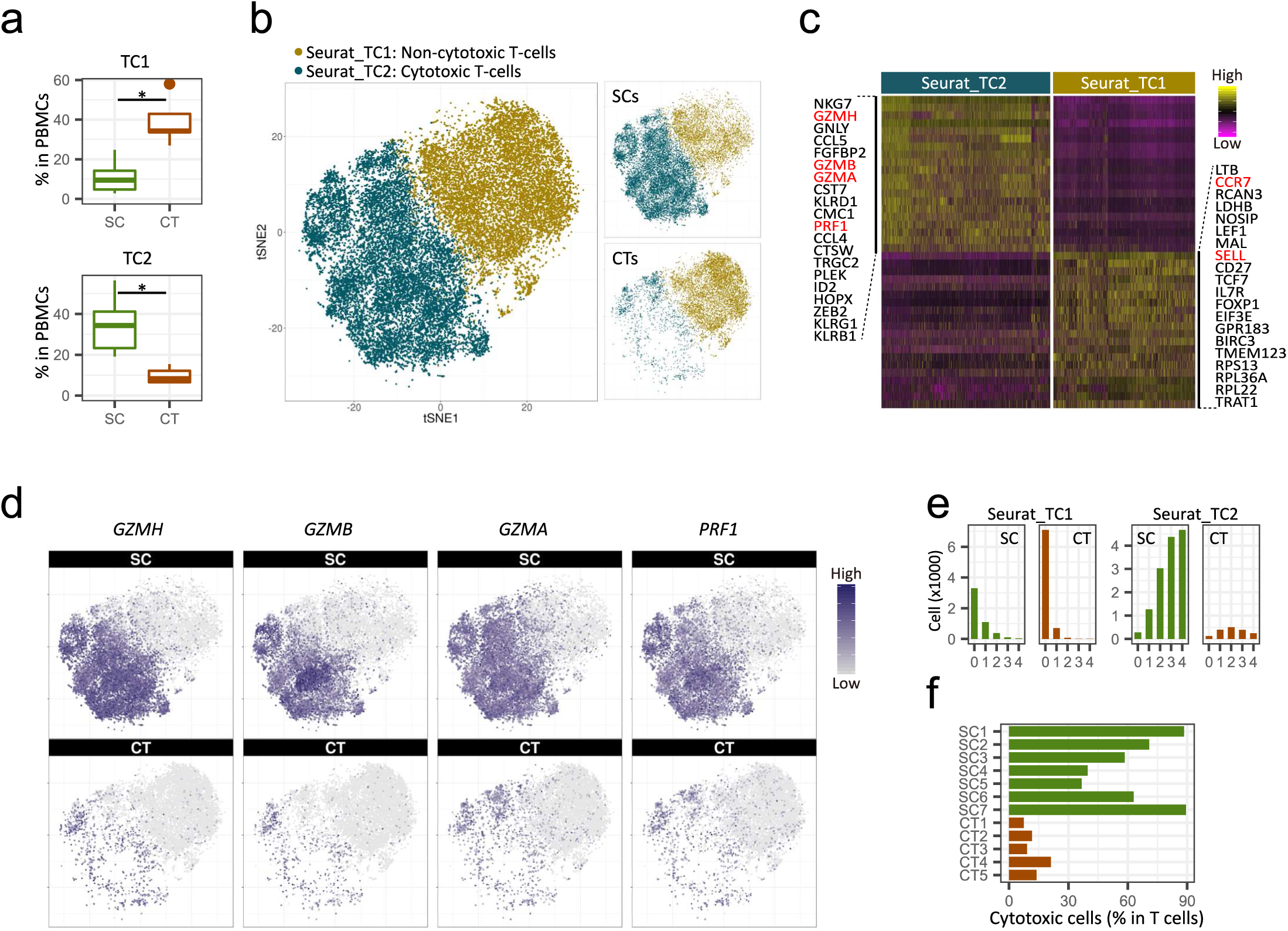
Expansion of cytotoxic T-cells in supercentenarians. **a.** Boxplots of percentages of TC1 and TC2 T-cells (defined by k-means clustering of single cell RNA-Seq data) in PBMCs of seven supercentenarians (SC1–SC7) and five controls (CT1–CT5). *, *P* < 0.05 (Wilcoxon rank sum test). **b.** Two-dimensional tSNE visualization of T cells using the Seurat R package. Different colors represent two clusters (Seurat_TC1 and Seurat_TC2), similar to the original TC1 and TC2 clusters. Right panels (top and bottom) show supercentenarians and controls, respectively. **c.** Top 20 genes significantly highly expressed in Seurat_TC2 (left) and Seurat_TC1 (right). Major cytotoxic effector genes and lymph node homing markers are shown in red. **d.** Expression of cytotoxic genes in supercentenarians (upper panels) and controls (lower panels); cell positions are from the tSNE plot in **b**. **e.** Number of detected genes out of four cytotoxic genes (*GZMH*, *GZMB*, *GZMA*, and *PRF1*) per cell. **f.** Percentage of cytotoxic T-cells (cells clustered in TC2) among the total T-cells.

### Expansion of cytotoxic CD4 T-cells in supercentenarians

In general, cytotoxic T-cells are CD8+ and non-cytotoxic helper T-cells are CD4+, with both being derived from double positive thymocytes (30). Therefore, a simple interpretation of our results is that there is an increase in CD8+ T-cells in supercentenarians. However, *CD8A* and *CD8B*, which encode the two components of CD8, were expressed only in a subset of cytotoxic T-cells, whereas *CD4* and *TRDC* (T-cell receptor delta constant) were expressed in the other subsets, suggesting the presence of three subsets of cytotoxic T-cells: CD8 CTLs (cytotoxic T-lymphocytes), CD4 CTLs, and γδ T-cells (Fig. 4a). To investigate cytotoxic T-cells other than CD8 CTLs, we manually defined CD4 CTLs and γδ T-cells based on ranges of *CD4*, *CD8*, and *TRDC* expression (Figs. 4a lower right and S4a). Previous studies reported that CD4 CTLs account for a tiny fraction of CD4+ T-cells in PBMCs (e.g., mean 2.2% in 64 healthy donors (31)). Here, the supercentenarians show significantly higher levels of CD4 CTLs (mean, 25.3% of total T-cells) than in the controls (mean, 2.8%) (*P* = 0.0025, Wilcoxon rank sum test), as well as higher levels of CD8 CTLs than in the controls (*P* = 0.0025), whereas the population of γδ T-cells was moderate in size and comparable to that in the controls (Figs. 4b and S4b). To validate the expansion of CD4 CTLs, we performed FACS analysis of six supercentenarians (SC1 and SC5– SC7 [studied above] and SC9 and SC10), one semi-supercentenarian (over 105 years old; SC8), and five controls (CT4 and CT5 [studied above] and CT6–CT8) (Fig. S1a) using antibodies against CD3, CD4, CD8, and GZMB. According to the CD4/CD8 staining profile (gated on CD3+), the T cells in the supercentenarians were not predominantly CD8+ T-cells (Figs. 4c and S4c). We then asked how many of the CD4+ T-cells retained in supercentenarians were cytotoxic by using the CD4/GZMB staining profile. Remarkably, CD4+GZMB+ T-cells were quite abundant in the supercentenarians, in which at least 10% (mean, 30.1%) of T cells are CD4 CTLs in all tested supercentenarian samples (n = 7) (Fig. 4d). The percentages of CD4 CTLs (CD4+GZMB+ T cells) in the total T-cell populations were significantly higher in the centenarians than in the controls (*P* = 0.018, Wilcoxon rank sum test) (Figs. 4e and S4d).

**Figure 4.**
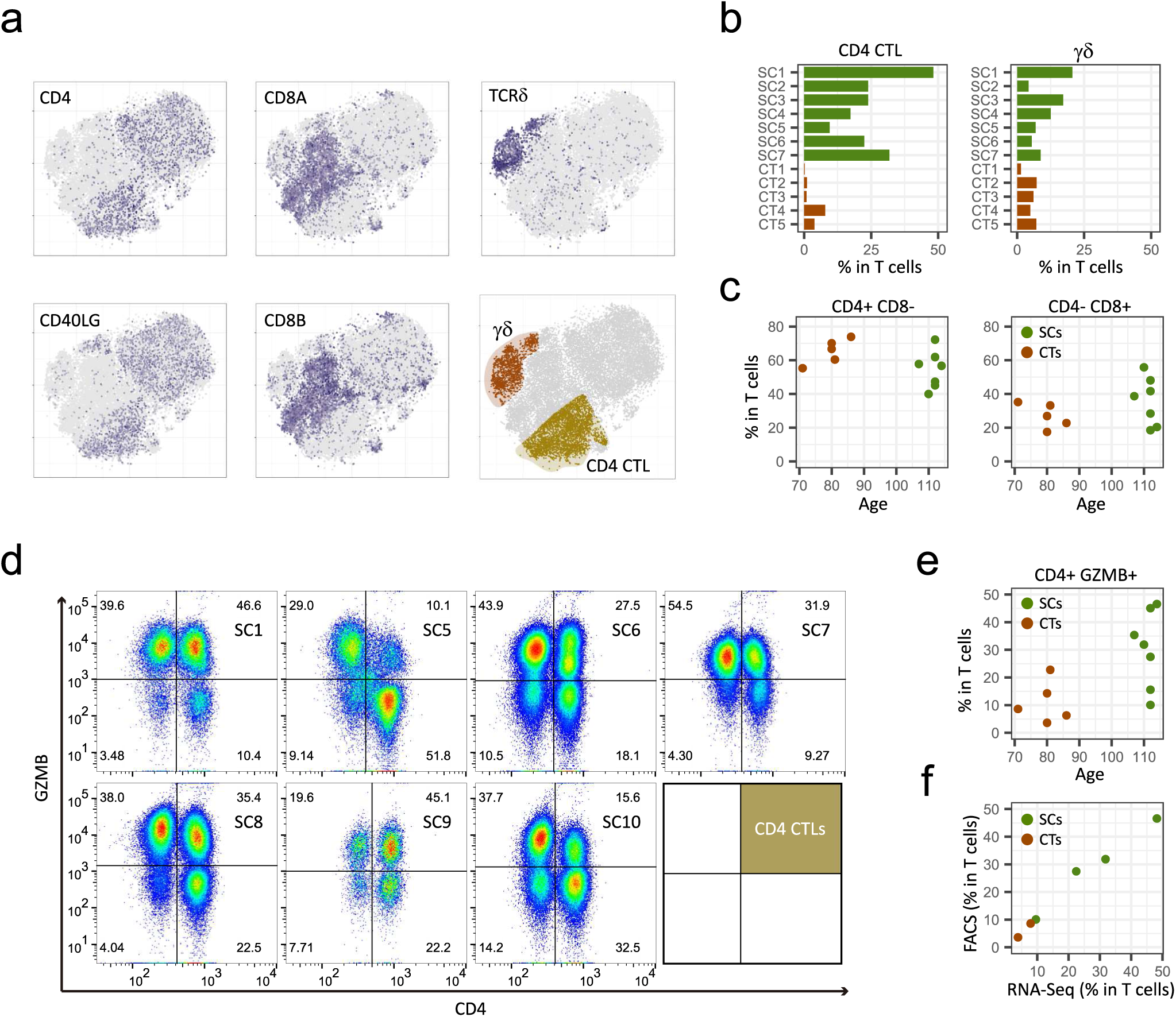
Expansion of cytotoxic CD4 T-cells in supercentenarians. **a.** Classification of cytotoxic T-cells into three subtypes—CD4 CTLs, CD8 CTLs, and γδ T-cells—was based on the expression of *CD4*, *CD8*, and *TRDC* (see also Fig. S4a) in T-cells of seven supercentenarians (SC1–SC7) and five controls (CT1–CT5); cell positions are from the tSNE plot in Fig. 3b. **b.** Percentages of CD4 CTLs and γδ T-cells among the total T-cells. **c.** Percentages of CD4+ T-cells and CD8+ T-cells in total T-cells. **d.** FACS profiles of six supercentenarians (SC1, SC5–7, and SC9) and one semi-supercentenarian (SC8). Cells gated on CD3+ were profiled using CD4 (x-axis) and GZMB (y-axis). Cells in upper right corners are CD4 CTLs. **e.** Percentages of CD4+ GZMB+ cells among the total T-cells of the six supercentenarians and one semi-supercentenarian listed in **d** and five controls (CT4, CT5, and CT6–CT8). **f.** Correlation between percentage of CD4 CTLs determined by RNA-Seq and FACS measurements. Each dot represents one donor, shown in green for supercentenarians (SC1, SC5–SC7) and red for controls (CT4, CT5).

Furthermore, GZMB+ cells were more abundant than GZMB− cells in both CD4 and CD8 T-cell populations in five out of seven tested (semi-)supercentenarians but none of the controls, indicating expansion of CD4 CTLs as well as CD8 CTLs (Fig. S4e). The percentages of CD4 CTLs correlated well between single-cell RNA-Seq and FACS analyses according to the comparison of the six commonly analyzed samples (four supercentenarians and two controls) (Fig. 4f). Thus, the high level of CD4 CTLs in supercentenarians was supported by two independent methods.

### Cell state transition of CD4 CTLs during T-cell differentiation

CD4 CTLs have been identified in differentiated T-cell subsets: i.e., effector memory (TEM) and effector memory re-expressing CD45RA (TEMRA) cells, which are often associated with a distinct surface phenotype including CCR7-, CD27-, CD28-, and CD11A+ (31, 32). To understand the CD4+GZMB+ T-cells in the context of differentiation, we constructed single-cell trajectories using the Monocle 2 (version 2.4.0) R package (33); all T cells in TC1 and TC2 were placed on these trajectories based on changes in their transcriptomes (Figs. 5a and S5a). Consistent with the clustering analyses, TC1 (the non-cytotoxic cluster) was mostly distributed throughout the early pseudotime, whereas TC2 (the cytotoxic cluster) was found mostly in later pseudotime, showing a clear temporal separation of the two (Fig. S5b). We then examined the transition of expression values along pseudotime for a panel of established marker genes associated with T-cell differentiation (29). As mentioned above, *CCR7* expression is a primary marker of central memory T-cells, and distinguishes them from effector memory T-cells. We observed rapid reduction of *CCR7* expression followed by the gradual loss of costimulatory molecules *CD27* and *CD28* (Fig. 5b) indicating that early pseudotime corresponds to naïve and central memory T-cells. The results also showed a gradual increase of expression of *GZMA*, *GZMB*, and *PRF1*, which encode cytotoxic molecules, as well as concordant patterns of expression of transcripts encoding adhesion and migration molecules (Figs. 5b and S5c) indicating progressive differentiation states of effector memory T-cells, corresponding to late pseudotime. One of the branches showed enriched expression of *FOXP3* and *IL2RA* (*CD25*), primary markers of regulatory T-cells (Figs. S5d and S5e). Altogether the backbone of pseudotime estimated by Monocle 2 recapitulated T-cell differentiation starting from naïve and central memory to terminally differentiated effector memory states with a branched trajectory of regulatory T-cell–like features. We examined the distributions of T cells along pseudotime separately for supercentenarians and controls. The T cells of the supercentenarians were clearly shifted toward more differentiated states compared with those of the controls (Fig. 5c): nearly 60% of T cells in the controls were placed in the earliest pseudotime corresponding to naïve and central memory T-cells, whereas T cells of supercentenarians were enriched in late pseudotime.

**Figure 5.**
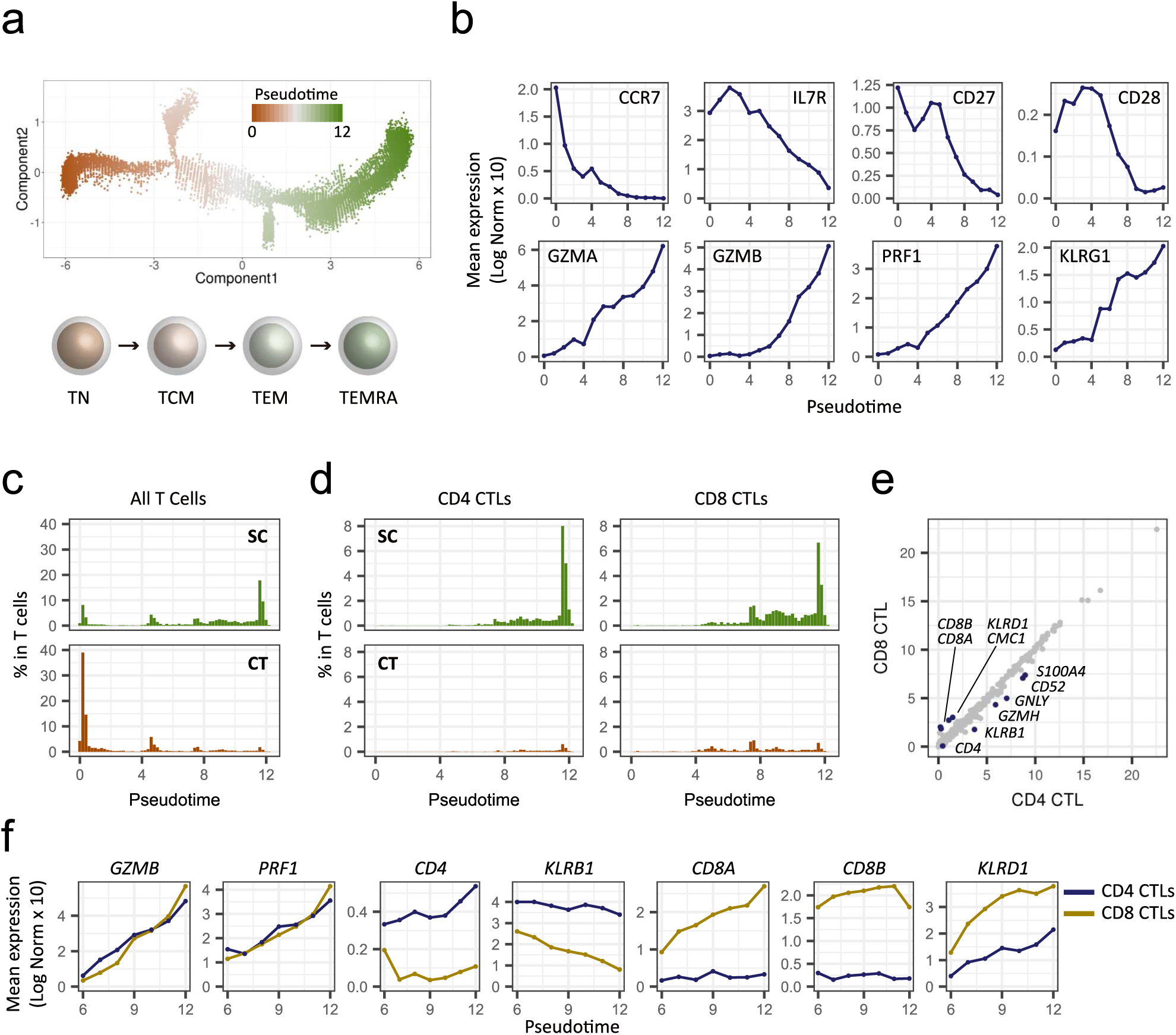
The differentiation state of T cells for seven supercentenarians (SC1–SC7) and five controls (CT1–CT5). **a.** Pseudotime trajectory of T cells estimated using Monocle 2. A continuous value from 0 to 12 was assigned to each cell as a pseudotime. The lower panel shows the general scheme of T-cell differentiation. TN, naïve; TCM, central memory; TEM, effector memory; and TEMRA, effector memory re-expressing CD45RA. **b.** Expression transition of differentiation-associated genes along the pseudotime. **c.** Percentages of T cells along the pseudotime for supercentenarians (SC) and controls (CT). **d.** Percentages of CD4 and CT8 CTLs among the total T-cells along the pseudotime. **e.** Correlation of gene expression between CD4 and CD8 CTLs. **f.** Expression transition of selected genes shown separately for CD4 and CD8 CTLs.

Next, we examined the distributions of CD4 CTLs (n = 5274) and CD8 CTLs (n = 7643), which were defined in Figures 4a and S4a. CD4 CTLs were distributed in the latter half of pseudotime in a similar way to CD8 CTLs (Figs. 5d and S5f), indicating a similar differentiation process despite fundamental functional differences between the two cell types. Indeed, mean expression values were highly correlated between CD4 and CD8 CTLs, with the exception of a small number of genes (Fig. 5e). The expression of four major cytotoxic genes *GZMA*, *GZMB*, *PRF1*, and *NKG7*, which are known to be abundant in CD4 CTLs (31, 34), increased along the latter half of pseudotime in a similar manner between CD4 and CD8 CTLs; however, the expression of two other major cytotoxic genes, *GZMH* and *GNLY*, showed slightly different patterns for CD4 and CD8 CTLs (Figs. 5f and S5g). Other exceptions were *KLRB1* and *KLRD1*, which encode two killer cell lectin-like receptors; at all time points, expression of these genes was higher in either CD4 or CD8 CTLs. In summary, we found a seemingly heterogeneous population of CD4 CTLs, which could be further categorized in pseudotime according to differentiation states. These differentiation states were characterized by progressive transcriptional changes, in a similar fashion to CD8 CTLs.

### Clonal expansion of CD4 CTLs

To explore the mechanism by which CD4 CTLs increased in supercentenarians, we performed an integrative analysis of the single-cell transcriptome and the T-cell receptor (TCR) repertoire. Firstly, we asked whether the high level of CD4 CTLs was reproducible at a different time point to that studied above. We re-collected fresh whole blood samples from two supercentenarians (SC1 and SC2) about 1.5 years after the first collection, and isolated CD4+ T-cells by negative selection (Fig. 6a). The single cell transcriptome profile generated using the Seurat R package confirmed high enrichment of T-cells, characterized by the expression of *CD3* genes (Figs. 6b and S6a). B-cells and CD14 monocytes were mostly depleted, whereas natural killer cells and erythroid cells were not completely depleted in the libraries (Fig. S6b). In the T-cell population, CD4+ T-cells were strongly enriched and CD8+ T-cells were depleted (Figs. 6c and S6c). We could recover transcripts encoding TCR alpha and beta chains in most of the T-cells, which were further clustered into two distinct cell types, based on the expression profiles (Figs. 6d and S6d). One of the clusters comprised CD4 CTLs, characterized by the co-expression of *GZMH*, *GZMA*, *GZMB*, *NKG7*, and *PRF1* as well as low expression of *SELL*, *CD27*, and *CCR7* (Figs. 6e, S6e, and S6f). CD4 CTLs accounted for about 62% (SC1) and 48% (SC2) of the CD4 T-cells in this analysis, which is consistent with the first sample collection from the same donor (Fig. 4b). This observation indicates that CD4 CTLs of supercentenarians are not transiently accumulated but persist in the blood for years.

**Figure 6.**
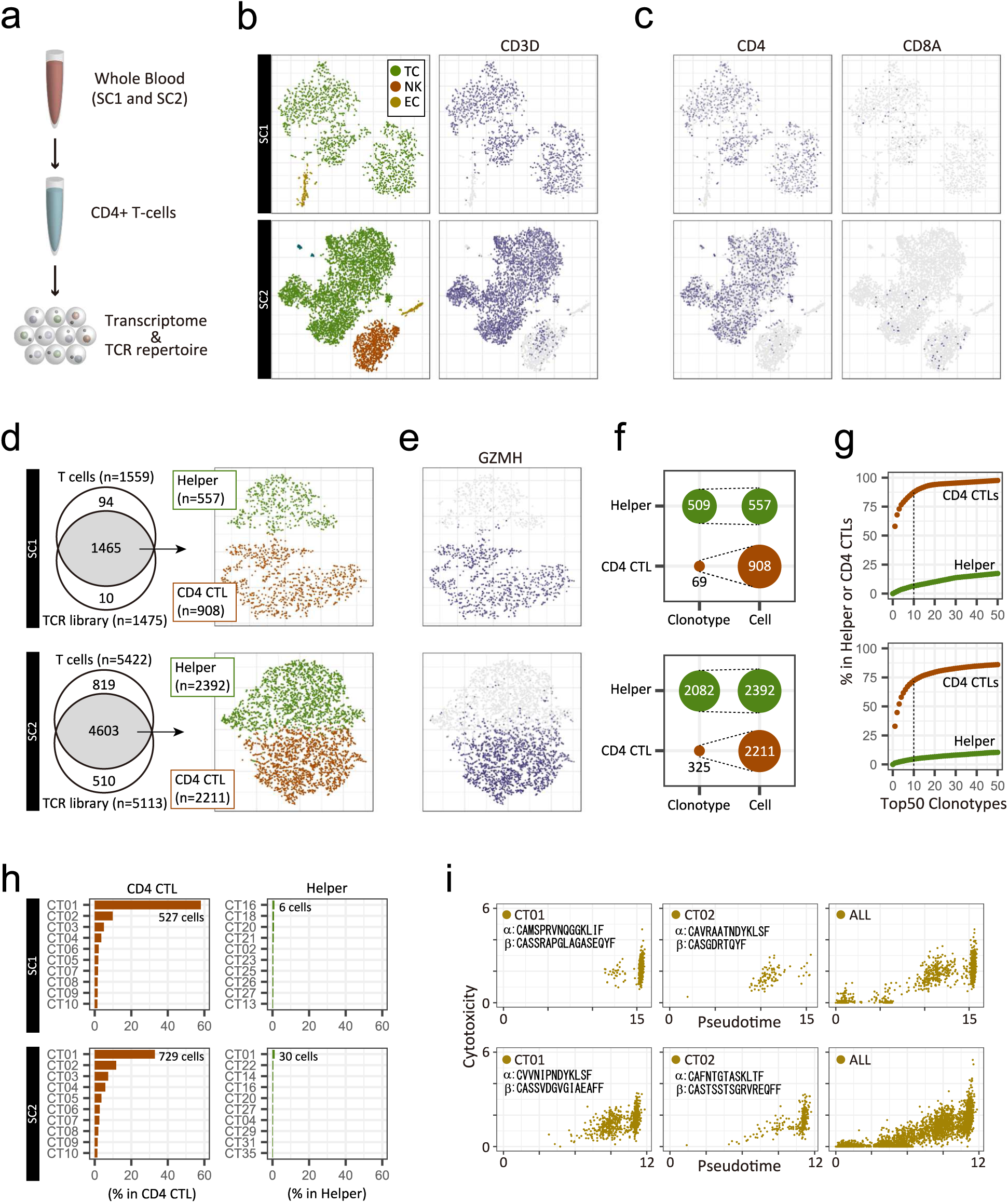
Single-cell transcriptome and TCR profiles of CD4+ T-cells for two supercentenarians (SC1, upper panels; and SC2, lower panels). **a.** Schematic representation of experiments for the single-cell transcriptome and TCR analysis. **b.** Two-dimensional tSNE visualization of three cell types (TC, T-cell; NK, natural killer cell; EC, erythroid cell), and CD3D expression (right). **c.** Expression of marker genes for CD4 and CD8 T-cells; cell positions are from the tSNE plot in **b**. T cells recovered in both transcriptome and TCR libraries. Recovered cells were clustered into helper T-cells and CD4 CTLs, shown in the tSNE plot. **e.** Expression of a marker gene for cytotoxic T-cells; cell positions are from the tSNE plot in **d**. **f.** Diversity of TCRs in helper T-cells and CD4 CTLs. **g.** Cumulative occupancy of the top 50 most abundant clonotypes. **h.** Occupancy of the top 10 most abundant clonotypes. **i.** Pseudotime and cytotoxicity of clonally expanded CD4 T-cells. Cytotoxicity values indicate the mean expression of five cytotoxic genes: *NKG7*, *GZMA*, *GZMB*, *GZMH*, and *PRF1*.

Secondly, we assessed the diversity of TCRs in CD4 CTLs and non-cytotoxic helper T-cells. We defined clonotypes based on CDR3 sequences of both TCR alpha and beta chains using the Cell Ranger analysis pipeline. We identified clonally expanded CD4 CTLs, which have only 69 clonotypes, among 908 cells in SC1; and 325 clonotypes among 2211 cells in SC2 (Fig. 6f). Moreover, the top 10 clonotypes occupied more than 70% of CD4 CTLs but less than 10% of helper T-cells (Fig. 6g). Both supercentenarians had one massively expanded clonotype, “CT01”, which accounted for 15% to 35% of the entire CD4 T-cell population (Fig. 6h), and had distinct combinations of TCR alpha and beta chains (TRAV12-3/TRAJ23 and TRBV3-1/TRBD2/TRBJ2-7 for SC1 and TRAV12-1/TRAJ20 and TRBV9/TRBD2/TRBJ1-1 for SC2). The cells of both “CT01” and “CT02” were mostly distributed in the CD4 CTL cluster with cytotoxic features (Fig. S6g) and were rarely placed in the CD4 helper T-cell cluster. The low TCR diversity of CD4 CTLs is in sharp contrast to helper T-cells in the same donor as well as younger controls, including publicly available CD4 and CD8 T-cells (Figs. 6h and S6h).

To understand the differentiation status of clonally expanded cells, we constructed a single-cell trajectory of CD4 T-cells using Monocle 2. The CD4 T-cells were distributed along pseudotime, following an increasingly differentiated trajectory, as evidenced by the marker gene expression patterns (Fig. S6i). As expected, at the late pseudotime, the top two expanded clonotypes CT01 and CT02 were enriched with highly expressed cytotoxic genes (Figs. 6i and S6i). Nevertheless, a subset of these cells was found in less differentiated states, indicating that a large number of CD4 T-cell clones with the same TCRs but at different levels of differentiation are circulating in the blood.

## Discussion

Here, we identified signatures of supercentenarians in circulating lymphocytes by using single-cell transcriptome analyses. In particular, CD4 CTLs were strongly expanded with distinct expression profiles including the activation of *GZMA*, *GZMB*, *GZMH*, *PRF1*, *NKG7* (*TIA-1*), *GNLY*, *CD40LG*, *KLRG1*, *KLRB1*, and *ITGAL* (*CD11A*) and the suppression of *CCR7*, *CD27*, *CD28*, and *IL7R* (Figs. 3d, 4a, 5b, and 5f). The results of single-cell TCR repertoire analysis of two supercentenarians suggest that the cell state transition of CD4 T-cells is at least partially explained by clonal expansion due to repeated stimulation with the same antigen. Here we discuss potential functions of CD4 CTLs in the late-stage of aging in terms of protective roles against tumor development and viral infections.

The primary function of CD4 T-cells, generally called helper T cells, is the regulation of immune responses using various cytokines, rather than direct elimination of target cells using cytotoxic molecules. Nevertheless, the presence of CD4 T-cells with cytotoxic features, namely CD4 CTLs, has been repeatedly reported in humans and mice (31, 32, 35). The reported fractions of CD4 CTLs are generally as low as a few percent of the total CD4 T-cells in healthy PBMCs (31, 36), whereas the size of the CD4 CTL fraction in the supercentenarians analyzed was on average 25% of T-cells, as measured by RNA-Seq and supported by the independent FACS measurements (Figs. 4b and 4e). More intriguingly, five out of seven supercentenarians analysed by FACS had more GZMB+ than GZMB− CD4 T-cells (Fig. 4d). The physiological role of the expanded CD4 CTLs remains unclear in humans, however a recent single-cell transcriptome study identified tumor-infiltrating CD4 CTLs in human hepatocellular carcinoma (23). In addition, several studies demonstrate that CD4 CTLs have the ability to directly kill tumor cells and eradicate established tumors in an MHC class II-dependent manner in mouse models (37, 38). Importantly, CD8 CTLs recognize class I MHC molecules present in nearly all cells. In contrast CD4 CTLs recognize class II MHC molecules, which are usually absent in normal non-immune cells, but present in a subset of tumor cells (39). This indicates that CD4 CTLs might contribute tumor immunity against established tumors, and may have an important role in immunosurveillance, helping to identify and remove incipient tumor cells abnormally activating class II MHC molecules.

Another potential function of CD4 CTLs is anti-viral immunity. A growing number of studies have demonstrated the direct cytotoxic activity, protective roles, and the associated induction of CD4 CTLs against various viruses such as dengue virus, influenza virus, hepatitis virus, CMV (cytomegalovirus), and HIV (human immunodeficiency virus) (40–44). Clonally expanded CD4 CTLs with virus-specific TCRs have been identified in dengue virus–positive donors (36). The association of CD4 CTLs with virus infection suggests that CD4 CTLs have accumulated in supercentenarians at least partially through clonal expansions triggered by repeated viral exposure. Although some important genes such as *CRTAM* and *ADGRG1* (*GPR56*) have been reported (32, 45), the exact molecular mechanism of the conversion from CD4 helper T-cells to CD4 CTLs is still unclear. Our transcriptome data show the striking similarity of gene expression and differentiation between CD4 CLTs and CD8 CTLs (Figs. 5d and 5e), suggesting that CD4 CTLs use the CD8 transcriptional program internally, while retaining CD4 expression on the cell surface. This agrees with the previous finding that CD4 helper T-cells can be reprogrammed into CD4 CTLs by the loss of ThPOK (also known as ZBTB7B), the master regulator of CD4/CD8 lineage commitment, with concomitant activation of CD8-lineage genes (46). The reinforcement of the cytotoxic ability by the conversion of CD4 T-cells in supercentenarians might be an adaptation to the late stage of aging, in which the immune system needs to eliminate abnormal or infected cells more frequently.

## Methods

### Human blood samples

All experiments using human samples in this study were approved by the Keio University School of Medicine Ethics Committee (approval number, 20021020) and the ethical review committee of RIKEN (approval number, H28-6). Fresh whole blood from supercentenarians, their offspring residing with them, and unrelated donors was collected in 2-ml tubes containing EDTA (ethylene diamine tetraacetic acid). PBMCs were isolated from whole blood within 8 h of sample collection by using SepMate-15 tubes (STEMCELL Technologies) with Ficoll-Paque Plus (GE Healthcare Life Sciences) according to the manufacturer’s instructions. Briefly, each blood sample was diluted with an equal volume of phosphate buffered saline plus 2% fetal bovine serum (FBS), added into a SepMate tube, and centrifuged at 1200 × *g* for 10 min at room temperature. Enriched mononuclear cells were washed with phosphate buffered saline plus 2% FBS and twice centrifuged at 300 × *g* for 8 min. Cell numbers and viability were measured using a Countess II Automated Cell Counter (Thermo Fisher Scientific).

### Single-cell library preparation

Single-cell libraries were prepared from freshly isolated PBMCs by using Chromium Single Cell 3ʹ v2 Reagent Kits (26). The cells and kit reagents were mixed with gel beads containing barcoded oligonucleotides (UMIs) and oligo dTs (used for reverse transcription of polyadenylated RNAs) to form reaction vesicles called GEMs (Gel Bead-in-Emulsions). The barcoded cDNAs in each GEM were pooled for PCR amplification, and adapter and sample indices were added. Single-cell libraries were sequenced with paired-end reads on the Illumina HiSeq 2500 platform, with mostly one sample per lane. The remaining PBMCs were suspended in CELLBANKER cryopreservation medium (ZENOAQ), and stored at −80°C.

### Single-cell data processing

The analysis pipelines in Cell Ranger version 2.1.0 were used for sequencing data processing. FASTQ files were generated using *cellranger mkfastq* with default parameters. Then, *cellranger count* was run with --transcriptome=refdata-cellranger-GRCh38-1.2.0 for each sample, in which reads had been mapped on the human genome (GRCh38/hg38) using STAR (version 2.5.1b) (47) and UMIs were counted for each gene. The outputs of *cellranger count* for individual samples were integrated using *cellranger aggr* with --normalize=mapped, in which read depths are normalized based on the confidently mapped reads. This command also runs principal component analysis (PCA), t-distributed stochastic neighbor embedding (tSNE), and k-means clustering algorithms to visualize clustered cells in two-dimensional space. The output of *cellranger aggr* was loaded into R by using an R package, Cell Ranger R Kit (version 2.0.0), developed by 10X Genomics (http://cf.10xgenomics.com/supp/cell-exp/rkit-install-2.0.0.R). Log-normalized expression values of all annotated genes were calculated using two functions, *normalize_barcode_sums_to_median* and *log_gene_bc_matrix*, implemented in the R package.

### Analysis of B-cell subsets

Cells categorized in the B-cell cluster by the k-means clustering were extracted and saved as a file using the *save_cellranger_matrix_h5* function in the R package Cell Ranger R Kit. This file was loaded into *cellranger reanalyze* to re-run PCA, tSNE, and k-means (k = 3) clustering algorithms. Wilcoxon rank sum test was applied to compare percentages of B-cells between the supercentenarians and controls using the wilcox.test function in R.

### Analysis of T-cell subsets

The Seurat R package (version 2.3.0) was used to analyze T-cell subsets (TC1 and TC2). The outputs of *cellranger count* were loaded using the *Read10X* function. Cells clustered in TC1 and TC2 by the Cell Ranger analysis pipelines were extracted, and principal components were calculated using *RunPCA* function. The first 16 principal components, based on the manual inspection of the elbow plot (*PCElbowPlot*), were used for cell clustering (using the *FindClusters* function with resolution 0.05) and tSNE visualization (using *RunTSNE*). Differentially expressed genes were identified using the *FindAllMarkers* function, and the top 20 genes were visualized in a heatmap using the *DoHeatmap* function. CD4 CTL, CD8 CTL, and γδ T-cell clusters were manually defined in the interactive mode of the t-SNE plot by using the *TSNEPlot* function with do.identify=TRUE based on the expression of marker genes. Wilcoxon rank sum test was applied to compare percentages of T-cell subtypes between the supercentenarians and controls using the wilcox.test function in R. *Pseudotime analysis*

Monocle 2 (version 2.4.0) was used to estimate a pseudo-temporal path of T-cell differentiation (33). Cells clustered in TC1 and TC2 by Cell Ranger analysis pipelines were loaded to create a Monocle object using the *newCellDataSet* function implemented in Monocle 2. The cells were ordered in pseudotime along a trajectory using *reduceDimension* with the DDRTree method and *orderCells* functions. Mean log-normalized expression values of selected marker genes were calculated for each bin from 0 to 12 pseudotime points.

### Antibodies and flow cytometric analysis

Cryopreserved PBMCs were thawed and suspended in FACS buffer (1× Hank’s Balanced Salt Solution with 2% FBS and 0.2% NaN3). Monoclonal antibodies specific for human CD3ε (UCHT1 and HIT3a), CD4 (RPA-T4), CD8 (RPA-T8), CD19 (HIB19), CD14 (M5E2), CD16 (B73.1), CD56 (B159), and GzmB (GB11) were purchased from BD Pharmingen. Cell numbers were counted using a Countess II Automated Cell Counter. For intracellular staining, cells were fixed and permeabilized with IntraPrep Permeabilization Reagent (Beckman Coulter) according to the manufacturer’s protocols. Cells were analyzed using FACSAria III and FACSAria SORP cell sorters (BD Biosciences) with FlowJo Software (version 10.4.2).

### Single-cell TCR analysis

RosetteSep Human CD4+ T Cell Enrichment Cocktail with SepMate-15 (STEMCELL Technologies) was used to remove non-CD4+ T-cells from fresh whole blood. A single-cell transcriptome library was prepared from the enriched CD4+ T-cells by using the Chromium Single Cell 5ʹ Library Kit (10X Genomics) with 50 ng of cDNA amplified product. A single-cell TCR library was prepared using Chromium Single Cell V(D)J Enrichment Kits, Human (10X Genomics). The libraries were sequenced with paired-end 150-bp reads on the Illumina HiSeq 2500 platform. Analysis pipelines in Cell Ranger version 3.0.2 (updated version was used for the 5’ single-cell and TCR libraries from version 2.1.0 used for the 3’ single-cell libraries) were used for the sequencing data processing. TCR data were processed by running cellranger vdj with -- reference=refdata-cellranger-vdj-GRCh38-alts-ensembl-2.0.0 to assemble TCR alpha and beta chains and determine clonotypes. Transcriptome data were processed by running cellranger count with --transcriptome=refdata-cellranger-GRCh38-1.2.0. The Seurat R package (version 2.3.0) was used for cell clustering (FindClusters) and tSNE visualization (RunTSNE). Three control datasets of T-cell clonotypes analyzed by the same 10X Genomics kits were downloaded from the 10X Genomics web sites below (need a simple registration).

T-cells: http://cf.10xgenomics.com/samples/cell-vdj/3.0.0/vdj_v1_hs_pbmc2_t/vdj_v1_hs_pbmc2_t_clonotypes.csv

CD4 T-cells: http://cf.10xgenomics.com/samples/cell-vdj/2.2.0/vdj_v1_hs_cd4_t/vdj_v1_hs_cd4_t_clonotypes.csv

CD8 T-cells: http://cf.10xgenomics.com/samples/cell-vdj/2.2.0/vdj_v1_hs_cd8_t/vdj_v1_hs_cd8_t_clonotypes.csv

## Data availability

Raw UMI counts and normalized expression values for single-cell RNA-Seq are publicly available at http://gerg.gsc.riken.jp/SC2018/. Individual sequencing data will be available on request under the condition of approval of the ethics committee of Keio University and material transfer agreement.

## Acknowledgements

We would like to thank all the donors who participated in this study. We also thank RIKEN GeNAS for the sequencing of the single-cell libraries. This work was supported by a Research Grant from the Japanese Ministry of Education, Culture, Sports, Science and Technology (MEXT) to the RIKEN Center for Integrative Medical Sciences and Research Grants for Keio University Global Initiative Research Projects.

## Author contributions

KH, MV, GP, and PC contributed bioinformatics analyses and interpretation of data. TK, NH, HY, TT, YO, JWS, and AM contributed experiments and data production of the single-cell transcriptome. TI, YM, TS, HS, and IT contributed FACS analysis and interpretation. TS, HO, YA, and NH contributed recruitment of supercentenarians and management of human ethics samples. KH, TS, GP, AM, IT, YA, NH, and PC contributed to planning the study. NH and PC supervised the project.

---

**Figure S1.**
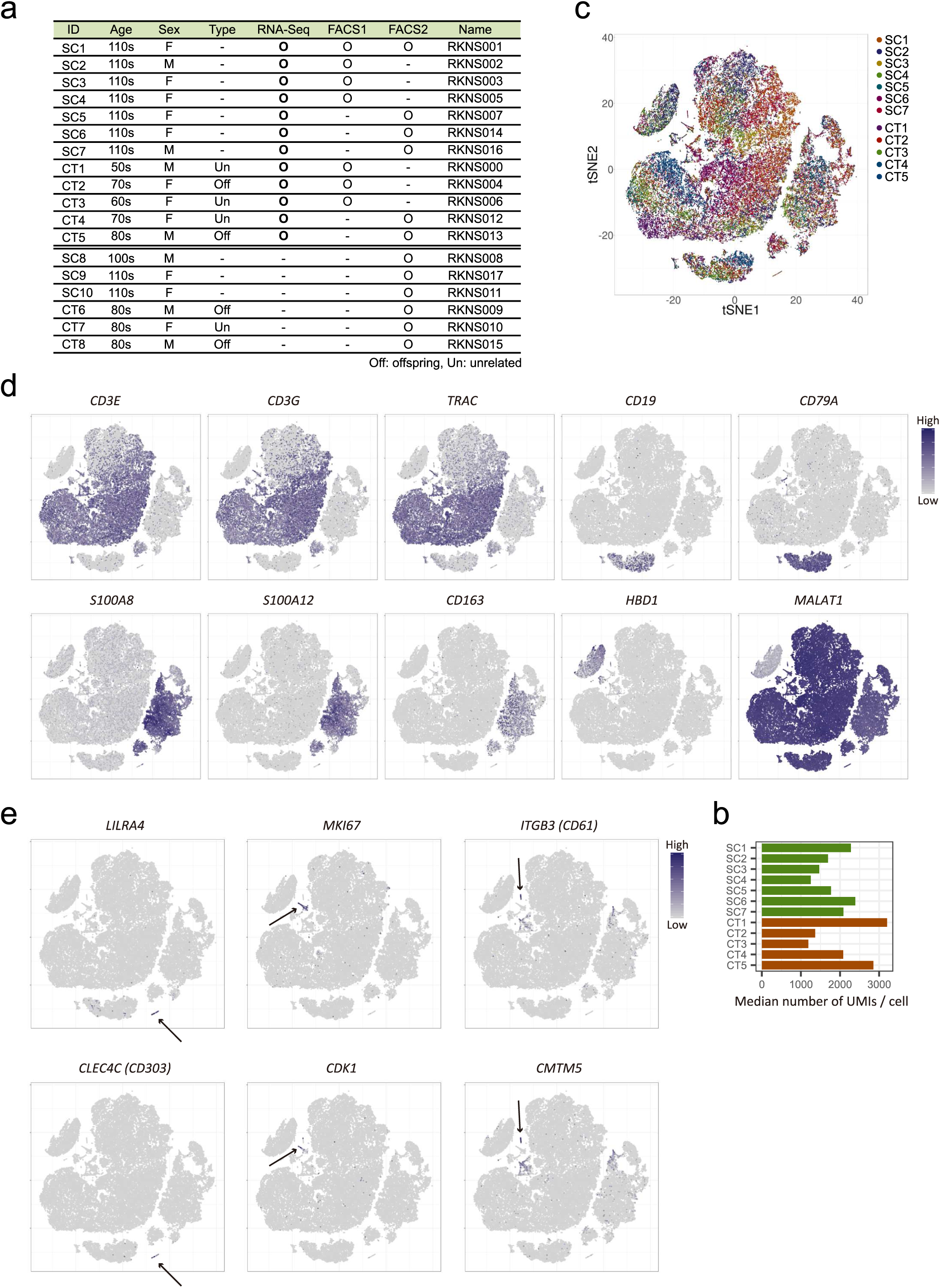
Single-cell transcriptome profile of PBMCs. **a.** Samples used for single-cell RNA-Seq and FACS analyses. FACS1 used antibodies against major cell-type markers: CD3, CD19, CD14, CD16, and NCAM1 (CD56), whereas FACS2 used antibodies against T-cell subtype markers: CD3, CD4, CD8, and GZMB. **b.** Median numbers of UMI counts per cell for each donor **c.** Two-dimensional tSNE visualization of 61,202 PBMCs. Different colors represent twelve donors. **d.** Expression of marker genes for cell-type markers and *MALAT1* as the highest expressed gene; cell positions are from the tSNE plots in Fig. 1c. **e.** Expression of marker genes used to define three small clusters by k-means clustering (i.e., MKI67+ proliferating cells [MKI], dendritic cells [DC], and megakaryocytes [MGK]).

**Figure S2.**
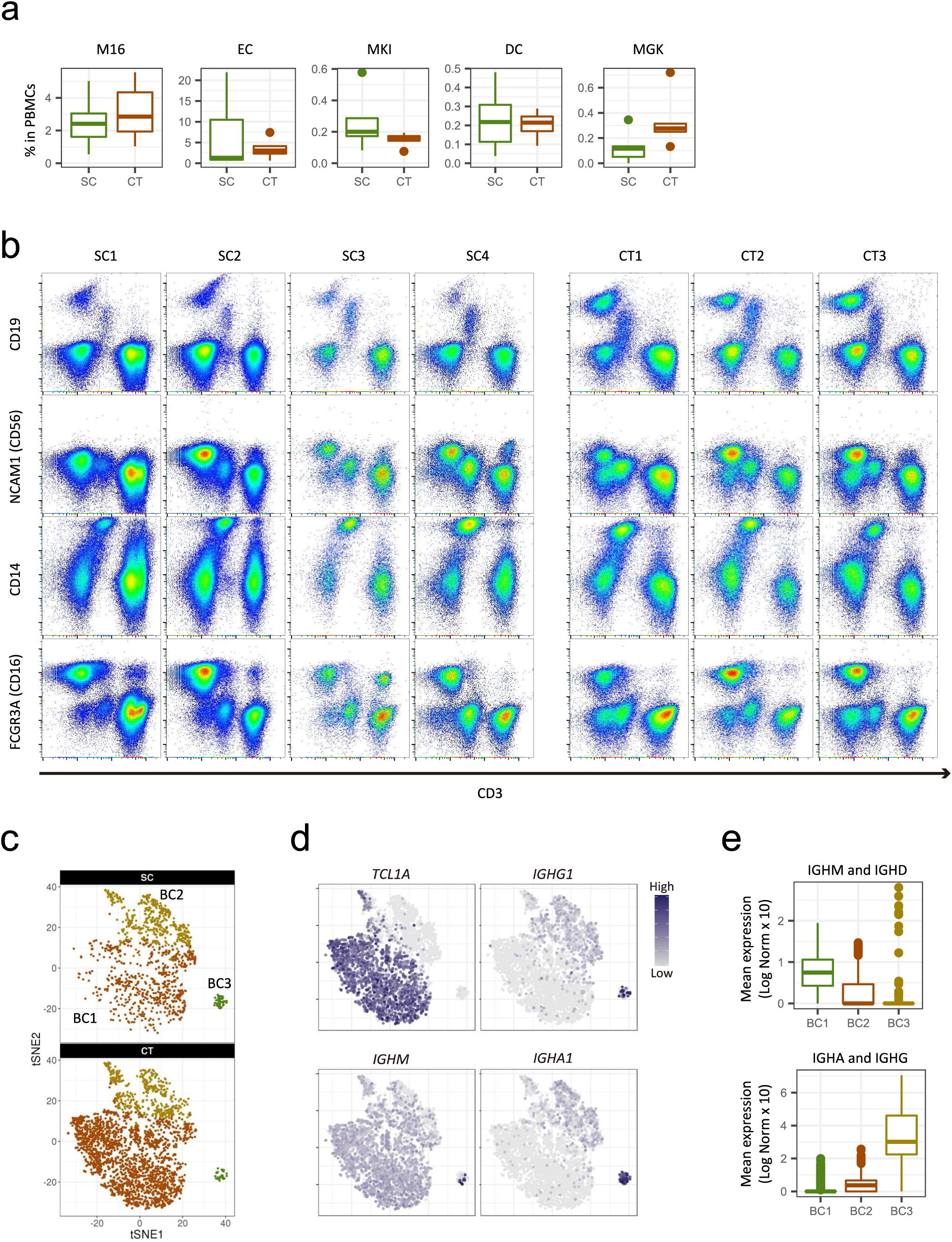
Reduction in the number of B cells in supercentenarians. **a.** Boxplots of the percentage of each indicated cell type in PBMCs. **b.** FACS plots for four supercentenarians (SC1–SC4) and three controls (CT1–CT3) profiled using CD3, CD19, NCAM1 (CD56), CD14, and FCGR3A (CD16). **c.** Two-dimensional tSNE visualization of B cells shown separately for supercentenarians (SC) and controls (CT). **d.** Expression of markers for B-cell subtypes; cell positions are from the tSNE plot in Fig. 2d. **e.** Expression of immunoglobulin heavy chains in each B-cell subtype. Upper panel shows total expression of IGHM and IGHD, which are used before the class switch. Lower panel shows total expression of IGHA1, IGHA2, IGHG1, IGHG2, IGHG3, and IGHG4, which are used after the class switch.

**Figure S3.**
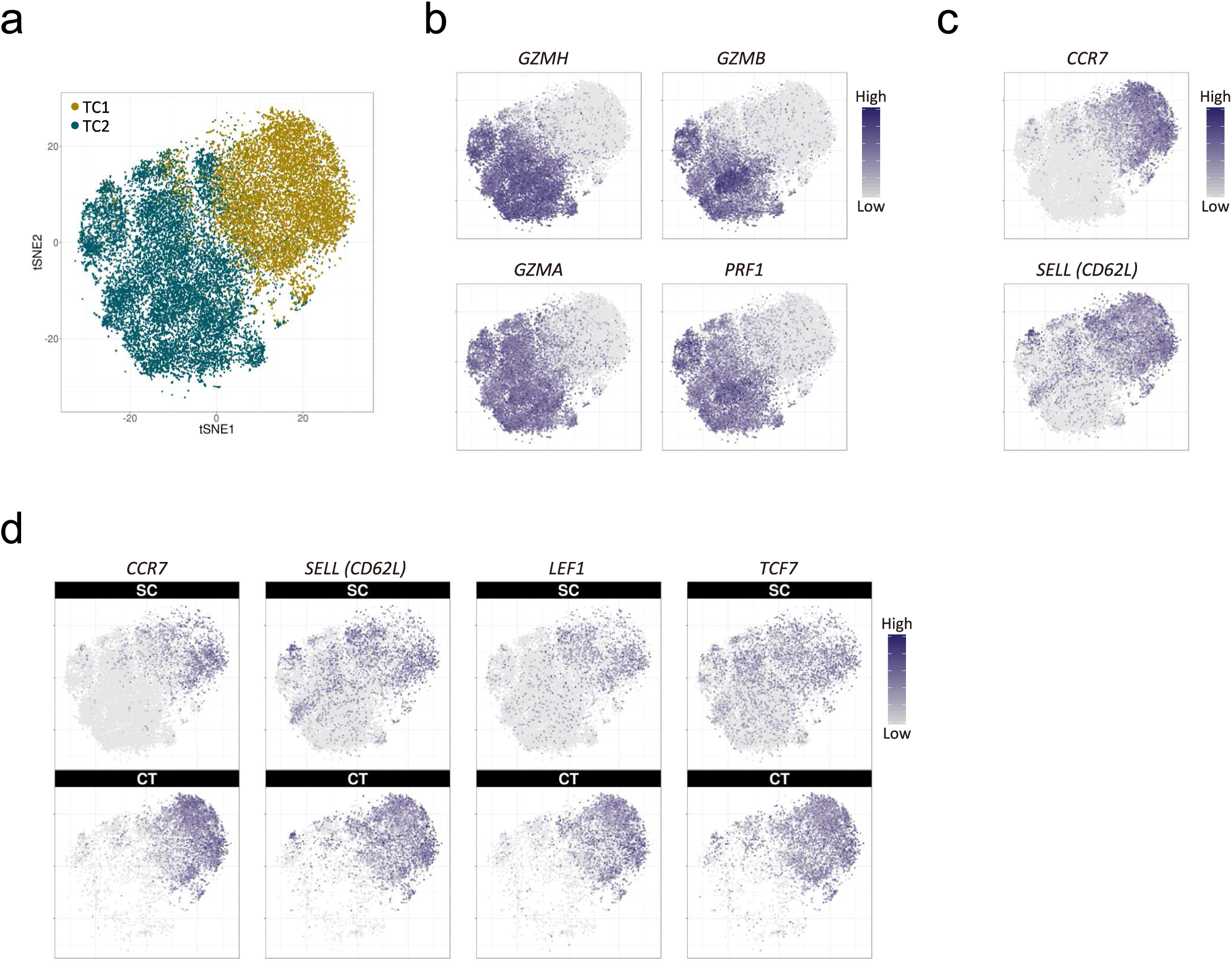
Expansion of cytotoxic T-cell populations in supercentenarians. **a.** Two-dimensional tSNE visualization of T cells using the Seurat R package. Different colors represent the original TC1 and TC2 clusters; 86.3% of the original TC1 is clustered into Seurat_TC1; 97.6% of the original TC2 is clustered into Seurat_TC2. **b–d.** Expression of cytotoxic genes and lymph node homing markers; cell positions are from the tSNE plot in **a.**

**Figure S4.**
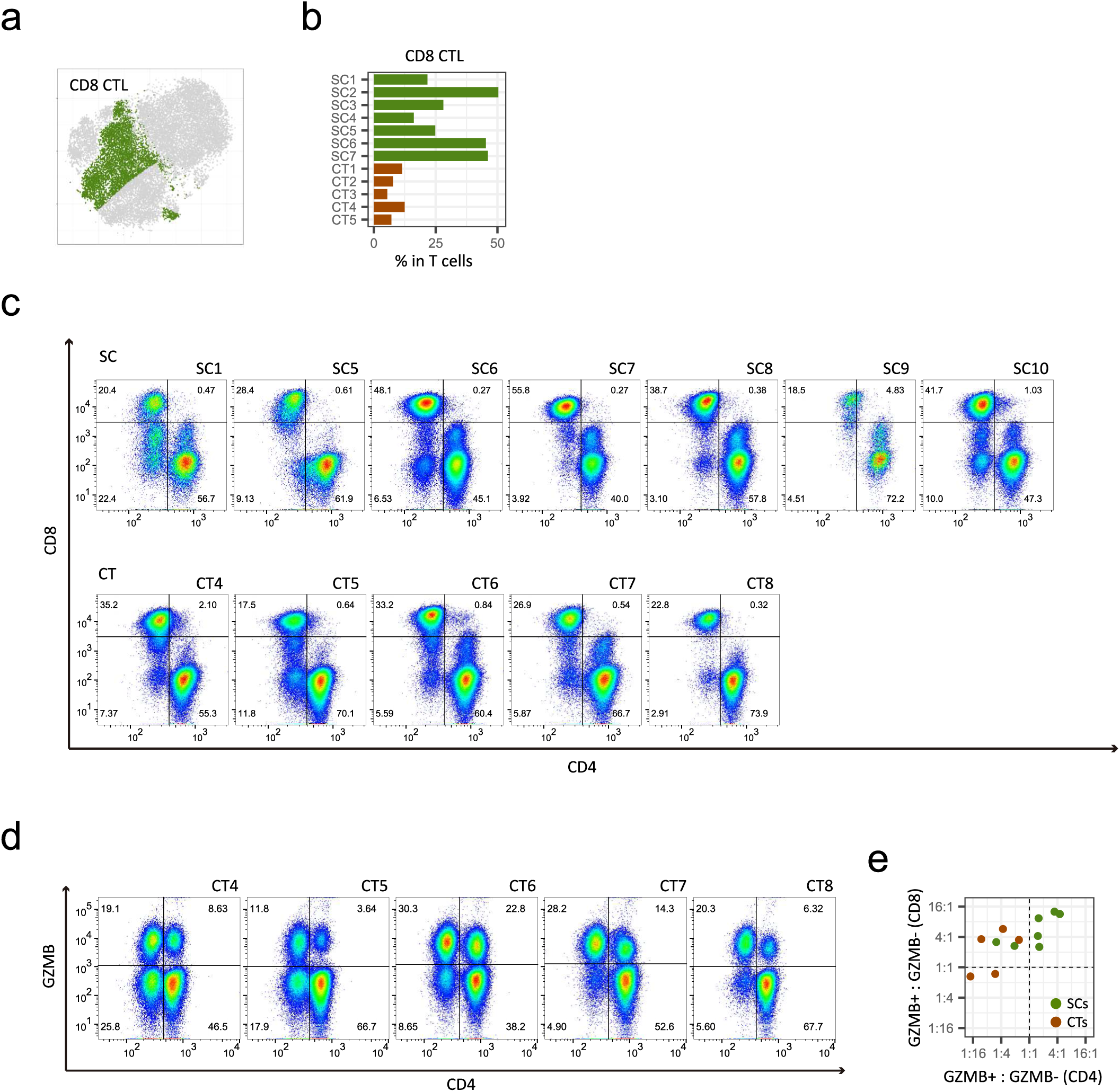
Expansion of cytotoxic CD4 T-cell populations in supercentenarians. **a.** CD8 CTLs defined based on the expression of *CD8A* and *CD8B*; cell positions are from the tSNE plot in Fig. 4a. **b.** Percentages of CD8 T-cells among the total T-cells. **c.** FACS profiles of supercentenarians (SC1, SC5–SC7, SC9, and SC10) and one semi-supercentenarian (SC8), and controls (CT4–CT8). Cells gated on CD3+ were profiled using CD4 (x-axis) and CD8 (y-axis). Cells in lower right and upper left corners are classified as CD4 and CD8 T-cells, respectively. **d.** FACS profiles of five controls (CT4–CT8). Cells gated on CD3+ were profiled using CD4 (x-axis) and GZMB (y-axis). Ratio between the percentage of GZMB+ and GZMB− cells in CD4 (x-axis) and CD8 (y-axis) T-cells. Ratio “1:1” indicates that the percentage of GZMB+ cells equals that of GZMB− cells.

**Figure S5.**
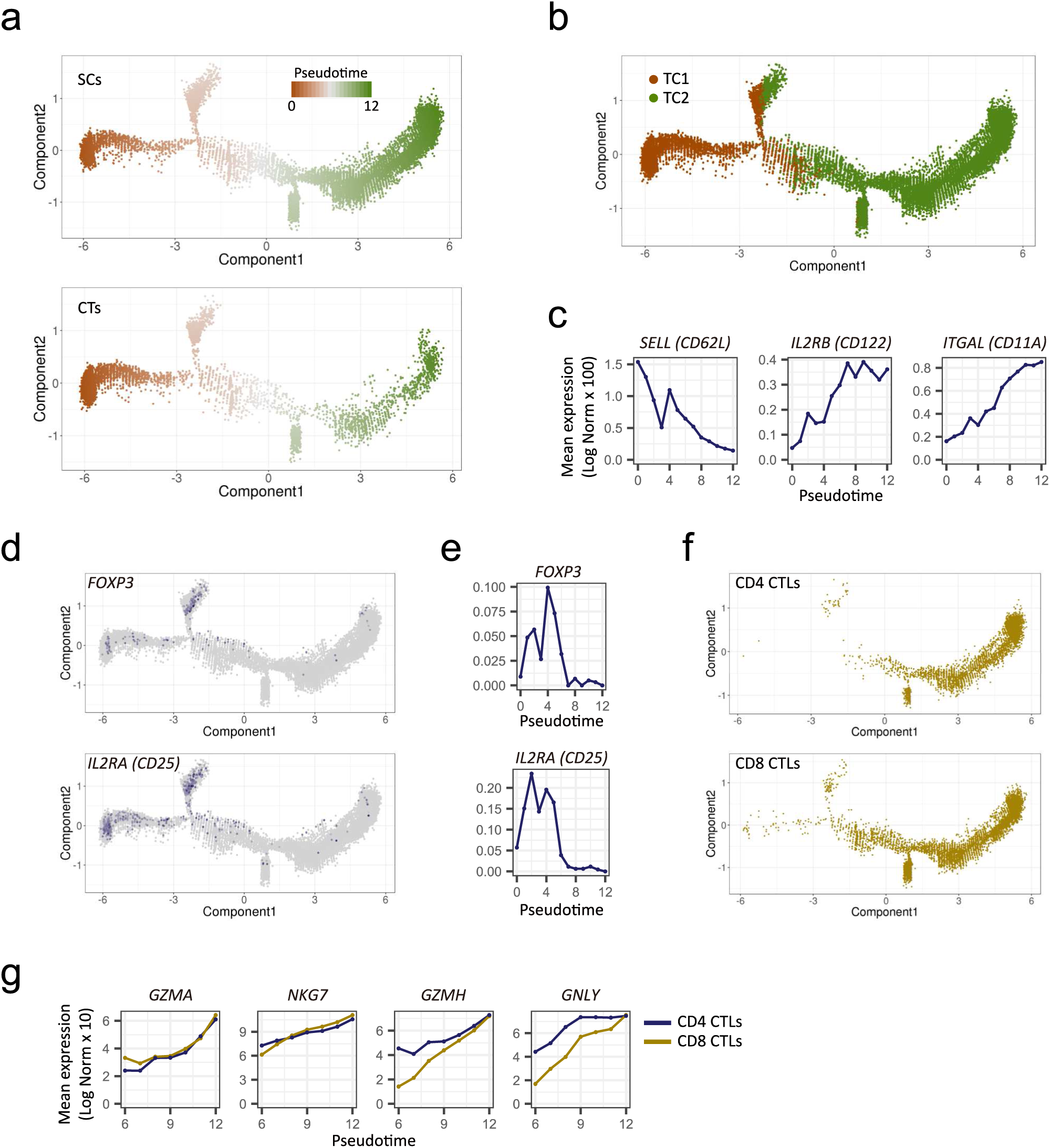
The differentiation state of T cells in supercentenarians. **a.** Pseudotime trajectory of T cells, shown separately for supercentenarians (SC) and controls (CT). **b.** Pseudotime trajectory of T cells colored by TC1 and TC2. **c.** Expression transition of differentiation-associated genes. **d.** Expression of *FOXP3* and *IL2RA* (CD25) mapped in pseudotime. **e.** Expression transition of *FOXP3* and *IL2RA* (CD25) along the pseudotime. **f.** Distribution of CD4 and CD8 CTLs mapped in pseudotime. **g.** Expression transition of selected genes shown separately for CD4 and CD8 CTLs.

**Figure S6.**
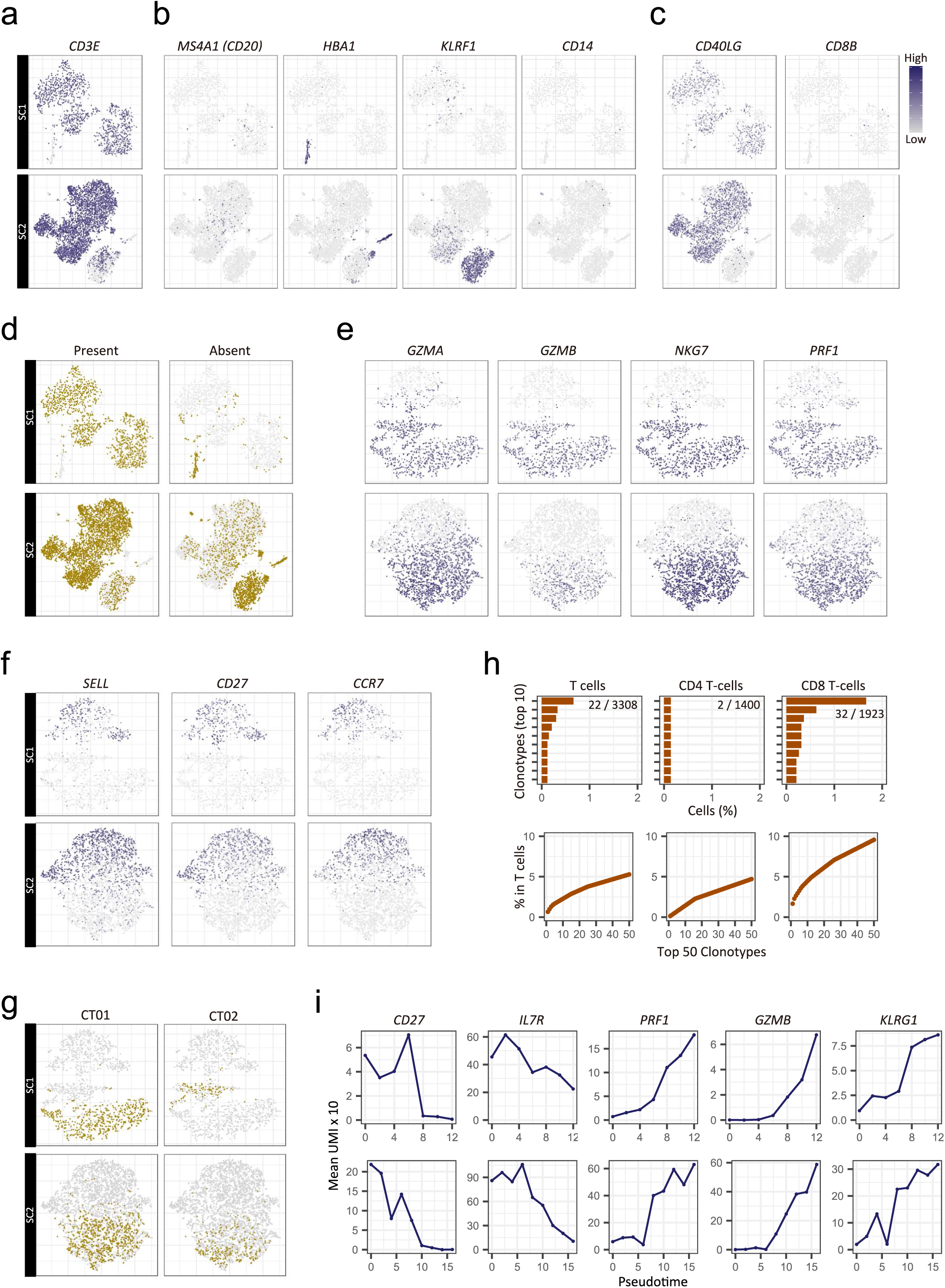
Single-cell 5ʹ transcriptome and TCR profiles for two supercentenarians (SC1, upper panels and SC2, lower panels). **a.** Expression of a marker gene for T-cells. **b.** Expression of marker genes for B-cells (*MS4A1*), erythrocytes (*HBA1*), NK cells (*KLRF1*), and monocytes (*CD14*). **c.** Expression of marker genes for CD4 T-cells (*CD40LG*) and CD8 T-cells (*CD8B*). **d.** Presence or absence of cells in TCR libraries. **e.** Expression of marker genes for cytotoxic T-cells. **f.** Expression of marker genes for T-cell differentiation. **g.** Distribution of the top 2 most abundant clonotypes, CD01 and CT02, on the t-SNE map. **h.** Diversity of TCRs in publicly available datasets released by another group. **i.** Transient expression of genes associated with T-cell differentiation.

## Reference

1. Young RD (2018) Validated Living Worldwide Supercentenarians, Living and Recently Deceased: February 2018. Rejuvenation Res 21(1):67–69.

2. Pavlidis N, Stanta G, & Audisio RA (2012) Cancer prevalence and mortality in centenarians: a systematic review. Crit Rev Oncol Hematol 83(1):145–152.

3. Willcox DC, et al. (2008) Life at the extreme limit: phenotypic characteristics of supercentenarians in Okinawa. J Gerontol A Biol Sci Med Sci 63(11):1201–1208.

4. Evert J, Lawler E, Bogan H, & Perls T (2003) Morbidity profiles of centenarians: survivors, delayers, and escapers. J Gerontol A Biol Sci Med Sci 58(3):232–237.

5. Arai Y, et al. (2014) Physical independence and mortality at the extreme limit of life span: supercentenarians study in Japan. J Gerontol A Biol Sci Med Sci 69(4):486–494.

6. Schoenhofen EA, et al. (2006) Characteristics of 32 supercentenarians. J Am Geriatr Soc 54(8):1237–1240.

7. Andersen SL, Sebastiani P, Dworkis DA, Feldman L, & Perls TT (2012) Health span approximates life span among many supercentenarians: compression of morbidity at the approximate limit of life span. J Gerontol A Biol Sci Med Sci 67(4):395–405.

8. Deeks SG (2011) HIV infection, inflammation, immunosenescence, and aging. Annu Rev Med 62:141–155.

9. Aw D, Silva AB, & Palmer DB (2007) Immunosenescence: emerging challenges for an ageing population. Immunology 120(4):435–446.

10. Arai Y, et al. (2015) Inflammation, But Not Telomere Length, Predicts Successful Ageing at Extreme Old Age: A Longitudinal Study of Semi-supercentenarians. EBioMedicine 2(10):1549–1558.

11. Chinn IK, Blackburn CC, Manley NR, & Sempowski GD (2012) Changes in primary lymphoid organs with aging. Semin Immunol 24(5):309–320.

12. Pang WW, et al. (2011) Human bone marrow hematopoietic stem cells are increased in frequency and myeloid-biased with age. Proc Natl Acad Sci U S A 108(50):20012–20017.

13. Rossi DJ, et al. (2005) Cell intrinsic alterations underlie hematopoietic stem cell aging. Proc Natl Acad Sci U S A 102(26):9194–9199.

14. Dunn-Walters DK & Ademokun AA (2010) B cell repertoire and ageing. Curr Opin Immunol 22(4):514–520.

15. Ademokun A, Wu YC, & Dunn-Walters D (2010) The ageing B cell population: composition and function. Biogerontology 11(2):125–137.

16. Sansoni P, et al. (1993) Lymphocyte subsets and natural killer cell activity in healthy old people and centenarians. Blood 82(9):2767–2773.

17. Fagnoni FF, et al. (2000) Shortage of circulating naive CD8(+) T cells provides new insights on immunodeficiency in aging. Blood 95(9):2860–2868.

18. Wertheimer AM, et al. (2014) Aging and cytomegalovirus infection differentially and jointly affect distinct circulating T cell subsets in humans. J Immunol 192(5):2143–2155.

19. Peters MJ, et al. (2015) The transcriptional landscape of age in human peripheral blood. Nat Commun 6:8570.

20. Stubbington MJT, Rozenblatt-Rosen O, Regev A, & Teichmann SA (2017) Single-cell transcriptomics to explore the immune system in health and disease. Science 358(6359):58–63.

21. Papalexi E & Satija R (2018) Single-cell RNA sequencing to explore immune cell heterogeneity. Nat Rev Immunol 18(1):35–45.

22. Enge M, et al. (2017) Single-Cell Analysis of Human Pancreas Reveals Transcriptional Signatures of Aging and Somatic Mutation Patterns. Cell 171(2):321–330 e314.

23. Zheng C, et al. (2017) Landscape of Infiltrating T Cells in Liver Cancer Revealed by Single-Cell Sequencing. Cell 169(7):1342–1356 e1316.

24. Chung W, et al. (2017) Single-cell RNA-seq enables comprehensive tumour and immune cell profiling in primary breast cancer. Nat Commun 8:15081.

25. La Manno G, et al. (2016) Molecular Diversity of Midbrain Development in Mouse, Human, and Stem Cells. Cell 167(2):566–580 e519.

26. Zheng GX, et al. (2017) Massively parallel digital transcriptional profiling of single cells. Nat Commun 8:14049.

27. Morbach H, Eichhorn EM, Liese JG, & Girschick HJ (2010) Reference values for B cell subpopulations from infancy to adulthood. Clin Exp Immunol 162(2):271–279.

28. Butler A, Hoffman P, Smibert P, Papalexi E, & Satija R (2018) Integrating single-cell transcriptomic data across different conditions, technologies, and species. Nat Biotechnol 36(5):411–420.

29. Mahnke YD, Brodie TM, Sallusto F, Roederer M, & Lugli E (2013) The who’s who of T-cell differentiation: human memory T-cell subsets. Eur J Immunol 43(11):2797–2809.

30. Taniuchi I & Ellmeier W (2011) Transcriptional and epigenetic regulation of CD4/CD8 lineage choice. Adv Immunol 110:71–110.

31. Appay V, et al. (2002) Characterization of CD4(+) CTLs ex vivo. J Immunol 168(11):5954–5958.

32. Tian Y, et al. (2017) Unique phenotypes and clonal expansions of human CD4 effector memory T cells re-expressing CD45RA. Nat Commun 8(1):1473.

33. Qiu X, et al. (2017) Single-cell mRNA quantification and differential analysis with Census. Nat Methods 14(3):309–315.

34. Zaunders JJ, et al. (2004) Identification of circulating antigen-specific CD4+ T lymphocytes with a CCR5+, cytotoxic phenotype in an HIV-1 long-term nonprogressor and in CMV infection. Blood 103(6):2238–2247.

35. Juno JA, et al. (2017) Cytotoxic CD4 T Cells-Friend or Foe during Viral Infection? Front Immunol 8:19.

36. Patil VS, et al. (2018) Precursors of human CD4(+) cytotoxic T lymphocytes identified by single-cell transcriptome analysis. Sci Immunol 3(19).

37. Quezada SA, et al. (2010) Tumor-reactive CD4(+) T cells develop cytotoxic activity and eradicate large established melanoma after transfer into lymphopenic hosts. J Exp Med 207(3):637–650.

38. Xie Y, et al. (2010) Naive tumor-specific CD4(+) T cells differentiated in vivo eradicate established melanoma. J Exp Med 207(3):651–667.

39. Haabeth OA, et al. (2014) How Do CD4(+) T Cells Detect and Eliminate Tumor Cells That Either Lack or Express MHC Class II Molecules? Front Immunol 5:174.

40. Weiskopf D, et al. (2015) Dengue virus infection elicits highly polarized CX3CR1+ cytotoxic CD4+ T cells associated with protective immunity. Proc Natl Acad Sci U S A 112(31):E4256–4263.

41. Brown DM, Lee S, Garcia-Hernandez Mde L, & Swain SL (2012) Multifunctional CD4 cells expressing gamma interferon and perforin mediate protection against lethal influenza virus infection. J Virol 86(12):6792–6803.

42. Aslan N, et al. (2006) Cytotoxic CD4 T cells in viral hepatitis. J Viral Hepat 13(8):505–514.

43. van Leeuwen EM, et al. (2004) Emergence of a CD4+CD28-granzyme B+, cytomegalovirus-specific T cell subset after recovery of primary cytomegalovirus infection. J Immunol 173(3):1834–1841.

44. Jellison ER, Kim SK, & Welsh RM (2005) Cutting edge: MHC class II-restricted killing in vivo during viral infection. J Immunol 174(2):614–618.

45. Takeuchi A, et al. (2016) CRTAM determines the CD4+ cytotoxic T lymphocyte lineage. J Exp Med 213(1):123–138.

46. Mucida D, et al. (2013) Transcriptional reprogramming of mature CD4(+) helper T cells generates distinct MHC class II-restricted cytotoxic T lymphocytes. Nat Immunol 14(3):281–289.

47. Dobin A, et al. (2013) STAR: ultrafast universal RNA-seq aligner. Bioinformatics 29(1):15–21.

